# Experimental Infection of greater white-toothed Shrews (*Crocidura russula*) with Borna disease virus 1: Insights into Viral Spread and Shedding

**DOI:** 10.1101/2024.07.30.605894

**Authors:** Daniel Nobach, Leif Raeder, Jana Müller, Sybille Herzog, Markus Eickmann, Christiane Herden

**Author notes:** DN, correspondence address: Chemical and Veterinary Analysis Agency Stuttgart (CVUAS), SchaflandstraBe 3/ 2, 70736 Fellbach, Germany, CH, correspondence address: FrankfurterstraBe 96, 35392 GieBen, Germany.

## Abstract

Numbers of human encephalitis cases caused by infection with Borna disease virus 1 (BoDV1) increase continuously in endemic areas. The reservoir host of BoDV1 is the bicoloured white-toothed shrew, albeit few naturally infected individuals of other shrew species have been detected. To establish a reliable experimental reservoir model, 15 greater white-toothed shrews were infected with a shrew-derived BoDV1 isolate by different inoculation routes (intracerebral, intranasal, oral, subcutaneous, intraperitoneal) and monitored up to 41 days. Except for the oral route all other animals (12/15) were successfully infected, and the majority of them displayed temporary reduced feed intake and loss of body weight but no inflammatory lesions. Infectious virus was isolated from 11/12 infected animals. Viral RNA was demonstrated by RT-qPCR in the central nervous system (CNS) and the majority of organs.

Immunohistochemistry demonstrated BoDV1 antigen in neurons and astrocytes in the CNS and peripheral nerves. High viral loads in the CNS and the spinal cord points towards spread from periphery to the CNS to enhance viral replication, and subsequent centrifugal spread to organs capable of secretion and excretions. In general, successful experimental BoDV1 infection of shrews proves their usefulness as animal model, enabling further studies on maintenance, transmission, pathogenesis, and risk assessment for human spill-over infections.

## Introduction

Diseases caused by pathogens originating from wildlife reservoirs are considered an emerging worldwide public health hazard [1,2]. In Germany, increasing numbers of human encephalitis cases, caused by infection with Borna disease virus 1 (BoDV1), have been discovered in ongoing studies and by retrospective analyses of cases with previous unknown aetiology [3–9].

BoDV1 is a negative-sensed single stranded RNA virus (order *Mononegavirales*, family *Bornaviridae*, genus *Orthobornavirus*) which has long been known to infect a wide range of vertebrates [10]. This includes primarily mammals and recently also humans, thereby substantiating its zoonotic capacity [3–5,7,9–12]. Most identified natural cases occurred in domestic livestock, such as horses, sheep, and South American camelids [10,13–16]. Natural infections are epidemiologically associated to the presence of its natural reservoir, the bicoloured white-toothed shrew (*Crocidura leucodon*) [6,17–20]. In accidental spillover hosts, BoDV1 infection can cause Borna disease, a severe neurological disorder with fatal meningoencephalitis and neurological symptoms, such as behavioural abnormalities, apathy, and ataxia [3–5,9–11,21]. Similar to animal spillover hosts, human BoDV1 infections lead to a severe and mostly fatal encephalitis [3–5,8,22]. The discovery of increasing numbers of acute human cases (>50 cases) demonstrate the impact of BoDV1 concerning public health [4,7,9,22–24]. However, the route of transmission of BoDV1 to humans and associated factors have not yet been fully elucidated despite case-control studies [25].

Epidemiological characteristics, like occurrence of natural infections in distinct endemic areas and higher incidence in certain years and months, have already pointed to a wildlife reservoir of BoDV1 [26–28]. So far, the identified main reservoir is the bicoloured white-toothed shrew (*Crocidura leucodon*) [17–19,29]. The presence of BoDV1 in other European shrews of the genus *Crocidura*, such as the greater white-toothed shrew (*Crocidura russula*) and the lesser white-toothed shrew (*Crocidura suaveolens*) have only recently been described, indicating either spillover events or potential additional wildlife reservoirs in some endemic areas [30].

The widespread tissue distribution and shedding of BoDV1 combined with the lack of corresponding gross and histological changes in naturally infected *C. leucodon* support their role as BoDV1 reservoir. *C. leucodon* harbours virus in nearly all organs, with the highest load in the central nervous system (CNS), and inflammation, especially encephalitis, as well as other CNS lesions or alterations in other organs are absent [18–20,29]. In contrast, BoDV1 is restricted to the CNS and peripheral nervous system in spillover hosts, where it causes a T-cell based immune-mediated inflammation, markedly non-purulent meningoencephalitis [3–5,7,11,22]. In persistently infected *C. leucodon*, virus presence and replication in organs such as kidney, skin, and salivary gland enables shedding via several routes like saliva, urine, and skin secretions [20]. Furthermore, clinical symptoms, as noted in spillover or dead-end hosts, are absent in *C. leucodon*. Persistently BoDV1-infected shrews neither show altered behaviour nor changes in feed intake or weight, despite continuous viral shedding, widespread virus distribution, and presence of virus-specific serum antibodies [20]. However, data on the transmission routes that ensure maintenance of BoDV1 in the reservoir population of *C. leucodon* are still lacking.

Infection research on BoDV1 infections has so far been hampered by the lack of suitable animal models for reservoir hosts. Various infection routes, the acute and chronic persistent phases of BoDV1 infections have already been studied extensively in spillover and experimental hosts, such as rats and mice [10,13,31,32]. In contrast, studies detailing the route of infection, viral spread and distribution, as well as potential occurrence of clinical signs and/or gross and histological lesions in the acute phase of BoDV1-infection of reservoir hosts are not available so far. Suitable shrew models for experimental studies did not yet exist.

Maintenance and breeding of *C. leucodon* as an insectivore species is more demanding than that of laboratory rodents and also in comparison to another shrew species of the genus Crocidura, the greater white-toothed shrew (*C. russula*). For *C. russula,* a successful captive breeding colony exists under laboratory conditions in our working group. Thus, this study aimed to establish and evaluate *C. russula* as a suitable and reliable shrew model for infection research on viruses, especially BoDV1.

*C. russula* can be found close to and in human habitats in extensive agricultural areas in Europe, including endemic areas of BoDV1, such as Germany and Switzerland [33–35]. The occurrence of *C. russula* and *C. leucodon* is considered partially sympatric with ecological separation, as the larger *C. russula* can displace the smaller *C. leucodon* [34,36,37]. The recent detection of BoDV1 in wild *C. russula* raises the question whether this species can serve as a BoDV1 reservoir in general [30]. In view of increasing numbers of human bornavirus infections and the unknown route of transmission, further investigations into the reservoir and host-virus-interactions are therefore urgently required. For this reason, the objectives of this study were as follows: firstly, to assess the feasibility of *C. russula* as shrew model for BoDV1 infections, and secondly, to characterize acute BoDV1 infections of *C. russula* with focus on different infection routes, clinical outcome, immune response, and viral shedding.

## Materials and Methods

### Ethics

The animal experiment was evaluated and approved by the ethics committee of the Regierungsprasidium GieBen (AZ G38/2018). All procedures were conducted in approved biosafety level 2 (BSL2) facilities according to animal welfare guidelines of the American Society of mammalogist [38] adopted to shrew husbandry.

### Animal Study

Greater white-toothed shrews (*Crocidura russula*) were bred at the Animal Facility of the Philipps-University Marburg licensed by the administrative district of Marburg-Biedenkopf (Az LRV FD 83.4.1-19c 20/21). Adult male and female shrews were infected via five different inoculation routes (oral, intranasal, subcutaneous, intraperitoneal, intracerebral) with a group size of three animals each. Infection was performed under general anaesthesia with isoflurane (induction 4% flow, maintenance 2% flow). All animals received at least 10 µl BoDV1 virus suspension (corresponding to virus dose of 6000 ID_50_) originally isolated from the skin of a naturally infected bicoloured white-toothed shrew [20]. The animals were weighed weekly and scored daily for clinical signs according to the adopted score sheet (Supplementary table 1), and feed intake was measured over a period of 41 days. Shrews were euthanized if moderate impairment of the animal’s welfare (as determined by scoring) was observed, or at the end of the study (day 41 post infectionem, p.i.) Additionally, oral swabs, skin swabs, and faeces were collected weekly under anaesthesia to monitor for virus shedding.

### Processing of Tissues

At necropsy, the following tissues were collected for histopathology and immunohistochemistry, and for RNA isolation and BoDV1-RT-PCR: cerebral cortex, hippocampus, olfactory bulb, brain stem, hippocampus, spinal cord (not for RNA isolation), brachial plexus, sciatic nerve, nasal conchae, parotid salivary gland, sublingual salivary gland, lung, heart, liver, spleen, kidney, urinary bladder, oesophagus, stomach, small intestine (duodenum), large intestine (colon), adrenal gland, genital tract (testis or ovary), skeletal muscle (hind leg), bone marrow (femur), dermal flank gland, and skin (abdomen) (overview in Table 1). For RNA isolation, tissues were taken aseptically and processed for RNA isolation with Qiazol and RNeasy Mini Kit according to manufacturer’s guide. Blood was collected postmortem and processed for detection of BoDV1-reactive serum antibodies. Additionally, a subset of tissues (hippocampus, lung, liver, kidney, urinary bladder, parotid gland, dermal flank gland) and blood were collected for infectivity tests as described previously [39]. Furthermore, sections of the above-mentioned tissues were fixed in 4% neutral-buffered formalin, embedded in paraffin and processed for histopathology and immunohistochemistry.

**Table 1.** Overview of collected tissues at necropsy.

| **BoDV1-RNA**<br>**(RT-qPCR)** | **serum**<br>**antibodies**<br>**(IIFT)** | **infectious virus**<br>**(virus isolation)** | **BoDV1 antigen**<br>**(IHC)** |
| --- | --- | --- | --- |
| brain (cerebral cortex) | blood | blood | brain (cerebral cortex) |
| brain (hippocampus) |  | brain (hippocampus) | brain (hippocampus) |
| brain (olfactory bulb) |  | lung | brain (olfactory bulb) |
| brain (brain stem) |  | liver | brain (brain stem) |
| brain (cerebellum) |  | kidney | brain (cerebellum) |
| peripheral nerve /<br>plexus brachialis |  | urinary bladder | peripheral nerve /<br>plexus brachialis |
| peripheral nerve /<br>ischiatric nerve |  | parotid salivary gland | peripheral nerve /<br>ischiatric nerve |
| parotid salivary gland |  | flank dermal gland | spinal cord (cervical) |
| sublingual salivary<br>gland |  |  | spinal cord (thoracal) |
| nasal conchae |  |  | spinal cord (lumbar) |
| lung |  |  |  |
| oesophagus |  |  | parotid salivary gland |
| stomach |  |  | sublingual salivary gland |
| small intestine |  |  | nasal conchae |
| large intestine |  |  | lung |
| liver |  |  | stomach |
| spleen |  |  | small intestine |
| bone marrow |  |  | large intestine |
| kidney |  |  | liver |
| urinary bladder |  |  | pancreas |
| genital tract |  |  | spleen |
| adrenal gland |  |  | kidney |
| skin |  |  | genital tract |
| flank dermal gland |  |  | adrenal gland |
| skeletal muscle |  |  | skin |
| heart |  |  | flank dermal gland |
|  |  |  | brown fat tissue |
|  |  |  | heart |

BoDV1, Borna disease virus 1. IIFT, indirect immunofluorescence test. IHC, immunohistochemistry.

RNA from oral swabs, skin swabs, and faeces was extracted with RNeasy Mini Kit according to manufacturer’s guide in an elution volume of 600 µl lysis buffer.

### Detection of BoDVI RNA by RT-qPCR

BoDV1 RNA was detected by quantitative RT-PCR (RT-qPCR) using BoDV1 primers and probe according to Schlottau et al. [3]. RT-qPCR runs were performed on Rotorgene Q (Qiagen) with SensiFAST Probe No-ROX One-Step Kit (Meridian Bioscience, formerly Bioline) according to manufacturer’s guide. The BoDV1 specific mix was combined with an internal beta-actin control mix to assess sample quality. A quantified BoDV1 standard curve as well as non-template controls were included in each RT-qPCR run.

### Detection of Bornavirus-reactive Serum Antibodies

Presence of bornavirus-reactive serum antibodies was investigated by indirect immunofluorescence assay as described elsewhere [18].

### Infectivity Test

Infectious BoDV1 was assessed by virus isolation with rabbit embryo brain cells followed by direct immunofluorescence test as described elsewhere [39,40]. Upon detection of contamination (e.g., mycotic overgrowth), tissues were subsequently excluded from virus isolation and infectivity testing.

### Histology and Detection of BoDVI Antigen by Immunohistochemistry

The above-mentioned tissues were examined histologically for the presence of inflammatory and/or degenerative lesions by two veterinary pathologists (DN, LR). Immunohistochemistry was performed applying the monoclonal antibody Bo18 detecting the BoDV1 nucleoprotein as described elsewhere [18]. Antigen distribution and number of infected cells was graded independently by two veterinary pathologists (DN, LR) according to the score established elsewhere [41]. An additional thorough interpretation of the antigen distribution within the spinal cord was performed by a board-certified pathologist (JM).

The experimental BoDV1 infection was considered successful if viral RNA and/or antigen was detected. Viral shedding was assessed positive in case of presence of infectious virus in swabs and/or tissues and/or by demonstration of viral RNA.

## Results

Experimental BoDV1 infection was successful via intranasal (3/3), subcutaneous (3/3), intraperitoneal (3/3) and intracerebral (3/3) inoculation of BoDV1 in greater white-toothed shrews, and not successful in orally inoculated animals (0/3). This indicates that experimental BoDV1 infection is easily possible in *C. russula* via different routes.

Most of the infected shrews displayed temporarily reduced feed intake and/or loss of body mass over the time course of the entire experiment (day 21 to 41, Figure 1). A first phase of reduced feed intake was observed in the 7 days immediately following BoDV1 infection performed under anaesthesia. However, no significant loss of body mass or other clinical signs, particularly neurological signs, were observed in any of the animals in these first days. From day 21 p.i. (dpi) onwards, a total of seven animals had to be euthanized due to reduced feed intake and body mass. Specifically, this involved one intranasally shrew on day 28 p.i. due to reduced feed intake and a loss of up to 23% of its starting body mass. In addition, two other shrews (one subcutaneously and one intraperitoneally infected animal) were removed from the experiment after day 28 p.i. for the same reasons, as were one intranasally infected and all intracerebrally infected animals from day 35 p.i. onwards (Table 2). None of the shrews showed any other obvious disturbance of general condition or clinical signs, other than the reduction of feed intake (40 % to 60 % of normal intake) and body mass.

**Figure 1.**
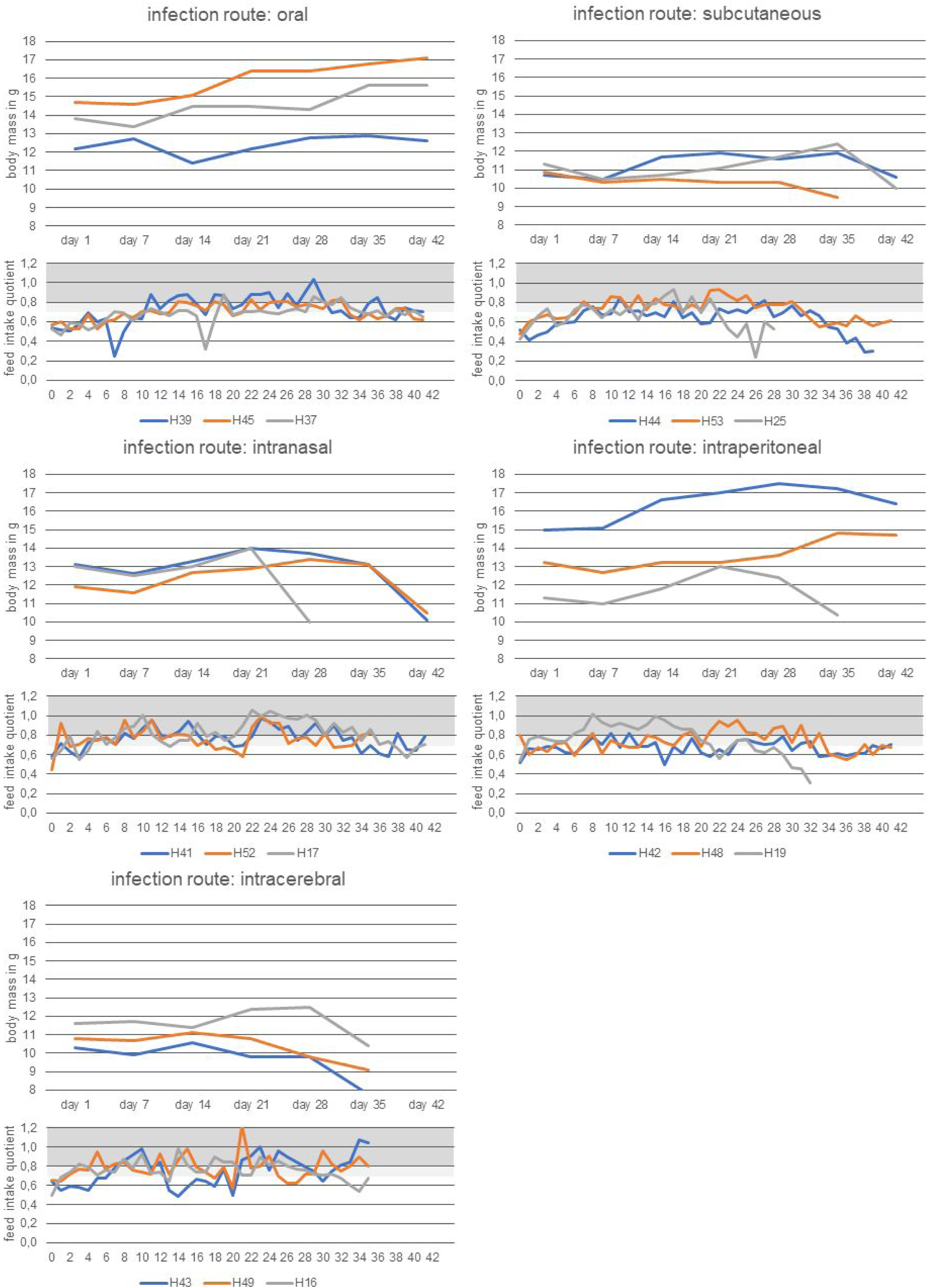
Development of body mass and feed intake during infection in shrews experimentally infected with BoDV1. Feed intake quotient is the quotient of daily feed intake and body mass. Dark grey area (feed intake quotient >0,8) is considered normal intake, light grey area (feed intake quotient between 0,7 and 0,8) is considered lightly reduced intake. BoDV1, Borna disease virus 1.

**Table 2.** Animals and characteristics of infection.

| **animal ID** | **sex** | **route of infection** | **end of trial<sup>a</sup>**<br>**[day p.i.]** | **titre of serum antibodies against BoDV1** | **detection of BoDV1 in brain<sup>b</sup>** |
| --- | --- | --- | --- | --- | --- |
| H39 | female | oral | 41 | < 1:10 | no |
| H45 | male | oral | 41 | < 1:10 | no |
| H37 | female | oral | 41 | < 1:10 | no |
| H44 | female | intranasal | 39 | 1:20480 | yes |
| H53 | male | intranasal | 41 | 1:40960 | yes |
| H25 | male | intranasal | 28 | 1:40960 | yes |
| H41 | female | subcutaneous | 41 | 1:40960 | yes |
| H52 | female | subcutaneous | 35 | 1:40960 | yes |
| H17 | male | subcutaneous | 41 | 1:40960 | yes |
| H42 | male | intraperitoneal | 41 | < 1:10 | no |
| H48 | male | intraperitoneal | 41 | 1:20480 | yes |
| H19 | female | intraperitoneal | 33 | 1:40960 | yes |
| H43 | female | intracerebral | 35 | 1:40960 | yes |
| H49 | female | intracerebral | 35 | 1:40960 | yes |
| H16 | male | intracerebral | 35 | 1:20480 | yes |

<sup>a</sup>End of trial before day 41 post infectionem (p.i.) was due to welfare score. <sup>b</sup>Detection of BoDV1 RNA via RT-qPCR and/or BoDV1-N-antigen via immunohistochemistry. Animal ID, animal identification number. BoDV1, Borna disease virus 1.

Infected shrews developed antibody titres of bornavirus-reactive serum antibodies ranging between 1:20480 – 1:40960 (Table 2) at the respective end point of the experiment without any significant difference between the animal groups.

In general, at the respective end point of the infection, viral RNA was consistently demonstrated by RT-qPCR in most organs, except for liver and spleen (Figure 2) in all infected cohorts (Supplementary table 2). Overall, the viral load was highest in the CNS of intracerebrally (i.c.) infected shrews with 2×10^10^ to 6×10^8^ copies on day 35 p.i. In the other groups, the highest viral load was also found in the CNS, ranging from 1×10^10^ to 1×10^7^ copies in the intranasal group, from 8×10^8^ to 1×10^6^ copies in the subcutaneous group, and 5×10^8^ to 1×10^6^ copies in the intraperitoneal group. Viral copy numbers mainly ranged between 2×10^8^ and 1×10^5^ copies in the parotid and sublingual salivary gland, with two highest values in the parotid gland of an intracerebrally infected animal (H49, 1,09×10^8^ copies), and in the sublingual salivary gland of an intranasally infected animal (h53, 2,56×10^8^ copies). In the gastrointestinal tract (oesophagus, stomach, small intestine, large intestine), viral copy numbers ranged between 1×10^8^ and 2×10^5^ copies, with the highest yields in stomach and small intestine of an intracerebrally infected animal (H16, 1,43×10^8^ and 1,39×10^8^ copies, respectively). In several animals, no viral RNA copies were detected in the urinary tract (kidney, urinary bladder). If detectable in this organ system, viral copy numbers ranged between 6×10^7^ and 3×10^4^ copies with the highest value in the urinary bladder of an intracerebrally infected animal (H49, 6,19×10^7^ copies). Concerning the respiratory tract (nasal conchae, lung), viral copy numbers were generally higher in the nasal conchae (1×10^9^ - 5×10^4^) in contrast to the lung (2×10^7^ - 3×10^4^ copies). The highest values were observed in the lung of an intracerebrally infected animal (H16, 2,42×10^7^ copies), and in the nasal conchae of an intranasally infected animal (H53, 1,3×10/\^9^ copies). In the intranasally infected group, viral load was high in the nasal conchae (4,1×10^5^ – 1,3×10^9^ copies) but only one animal had detectable viral load in the lung (H53, 8,21×10^6^ copies). In the skin and flank gland, viral copy numbers ranged mainly between 1×10^8^ and 7×10^5^ with consistent detection in the skin of all animals. The highest values were observed in an intracerebrally infected animal (H49, 1,25×10^8^ copies). In peripheral nerves (sciatic nerve, brachial plexus), viral copy numbers ranged mainly between 3×10^8^ and 1×10^5^ with the highest in the brachial plexus and sciatic nerve of an intracerebrally infected animal (H49, 3,0×10^8^ and 4,13×10^7^ copies, respectively). In the adrenal gland, the highest value was detected in a subcutaneously infected animal (H41, 1,43×10^8^ copies), but mainly ranged from7×10^7^ to 4×10^6^ copies in the other animals. In the genital tract (testis or uterus), viral copy numbers ranged from 4,29×10^7^ copies (H49, intracerebral infection) to 1,03×10^5^ copies (H52, subcutaneous infection) with detectable RNA yields in a total of six animals. In the bone marrow, viral copy numbers ranged between 1×10^8^ and 4×10^5^ copies with the highest value in an intracerebrally infected animal (H43, 1,19×10^8^ copies). In the heart, the highest value was observed in a subcutaneously infected animal (H41, 3,27×10^6^ copies), but for all the other animals, viral copy numbers ranged mainly between 3×10^6^ and 5×10^5^ copies. In the skeletal muscle, viral copy numbers ranged mainly between 4×10^7^ and 1×10^5^ copies with the highest in an intracerebrally infected animal (H49, 4,09×10^7^ copies). In the spleen, virus RNA was detected in a total of five animals (H16, H41, H48, H52, H53) with the highest copy number in an intraperitoneally infected animal (H48, 7,04×10^6^ copies). In the liver, no viral RNA was found in any infected shrew.

**Figure 2.**
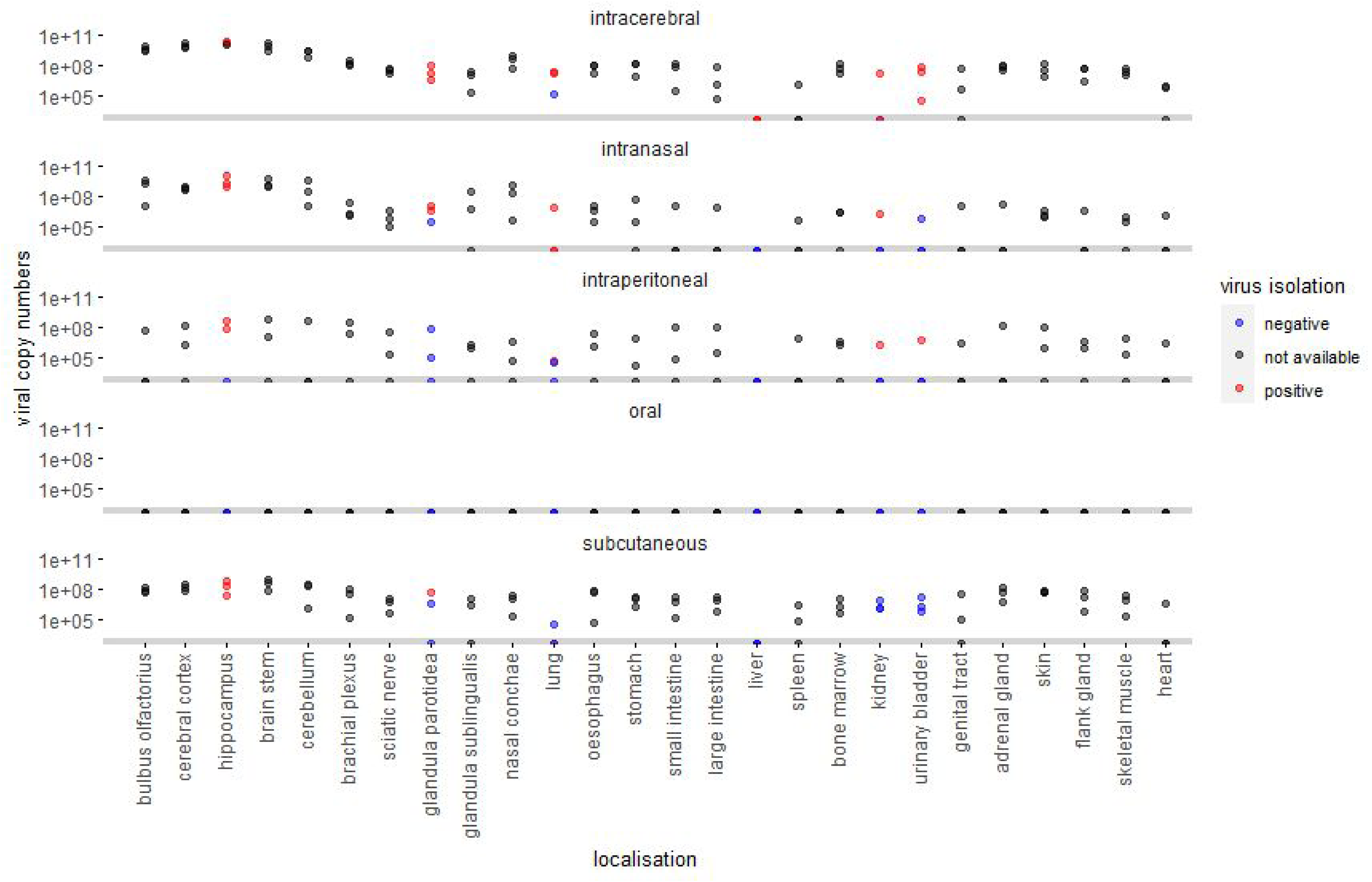
Overview of viral load in tissues and virus isolation in shrews experimentally infected with BoDV1 via different infection routes (intracerebral, intranasal, intraperitoneal, oral, and subcutaneous). The grey area depicts viral copy numbers below the lower limit (<1000 viral copies) and corresponds to Cq values above cycle 35. Blue and red dots represent unsuccessful and successful BoDV1 isolation, respectively. Grey dots represent tissue from which virus isolation was not performed or not available due to contamination (termed not available). BoDV1, Borna disease virus 1.

Concerning oral swabs, skin swabs, and faeces, BoDV1 RNA was first detected in a skin swab of a subcutaneously infected shrew at 35 dpi, followed by a positive skin swab of an intranasally infected shrew at 41 dpi, and detection in faeces of a subcutaneously infected shrew at 41 dpi. Additionally, infectious virus was detected in oral swabs of two intracerebrally infected shrews at 35 dpi (Figure 3).

**Figure 3.**
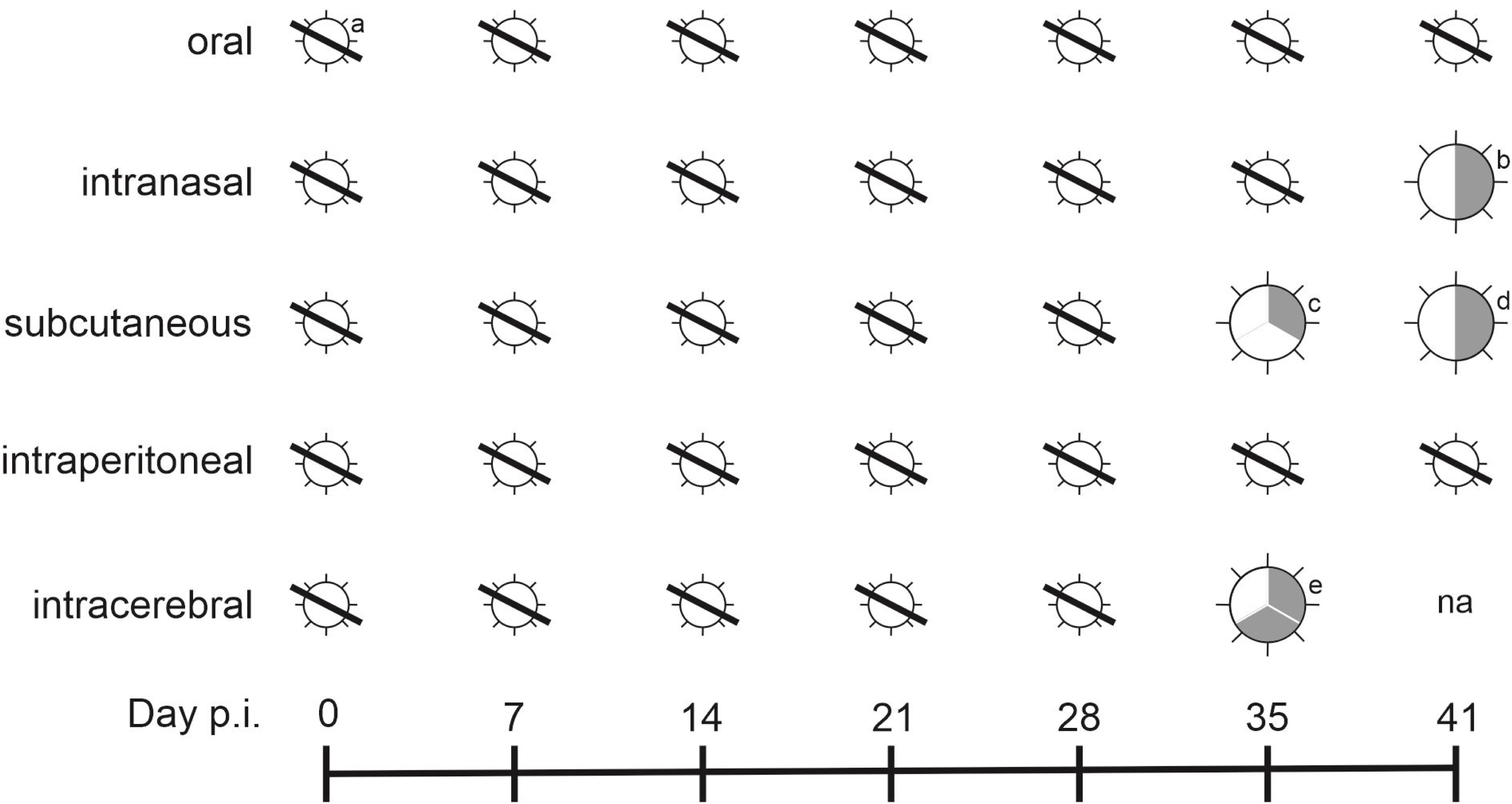
Schematic representation of detection of BoDV1-RNA and infectious virus in skin swabs, oral swabs and faeces. a) no detection, b) day 41, one of two intranasally infected shrews with RNA in skin swabs, no RNA or infectious virus in other swabs c) day 35, one of three subcutaneously infected shrews with RNA in skin swab, no RNA or infectious virus in other swabs, d) day 41, one of two subcutaneously infected shrews with RNA in faeces, no RNA or infectious virus in other swabs, e) day 35, two of three intracerebrally infected shrews with infectious virus in oral swabs, no RNA or infectious virus in other swabs, BoDV1, Borna disease virus 1. na, not available. p.i., post infectionem.

In general, infectious virus was consistently isolated from all successfully infected animals except one intraperitoneally infected animal. Isolation was successful from the CNS, and additionally from other organs such as lung, parotid salivary gland, urinary bladder, kidney, liver, and blood (Figure 2).

Euthanasia-associated lesions, such as alveolar oedema and haemorrhage, were observed in all animals on gross and histopathological examination. In contrast, none of the BoDV1-infected animals showed typical BoDV1-infection associated lesions described in the CNS of accidental hosts, such as inflammation or degeneration. Moreover, no gross or histological alterations were found in any other organ system examined.

BoDV1 antigen, demonstrated by immunohistochemistry, was predominantly found both in the cytoplasm and nucleus of neurons and astrocytes in the central and peripheral nervous system, including myenteric ganglia and chromaffin cells of the adrenal medulla (Table 3, Figure 4). The highest numbers of positive cells in the brain were found in intracerebrally and intranasally infected shrews. Both grey and white matter was affected, and in the hippocampus, antigen was found intracytoplasmic and intranuclear in neurons and astrocytes, as well as in the neuropil. In the cerebral cortex, antigen was present intracytoplasmic and intranuclear in neurons and astrocytes. In the brain stem, clusters of neurons and astrocytes displayed intracytoplasmic and intranuclear antigen. Few cerebellar Purkinje cells, including their axons, harboured intracytoplasmic and intranuclear antigen, and the molecular layer was spared. Notably, one subcutaneously infected animal (H52) displayed no positive cells in the cerebellum while antigen was found in neurons and astrocytes in the cerebral cortex, hippocampus and brain stem. In the olfactory bulb, antigen was found intracytoplasmic and intranuclear in neurons and astrocytes in all successfully BoDV1-infected groups, but not necessarily in every individual animal of the group. Thus, all intranasally and all intracerebrally infected animals displayed antigen-positive cells in this region, whereas BoDV1-antigen was not consistently detected in each of the subcutaneously and intraperitoneally infected animals.

**Figure 4.**
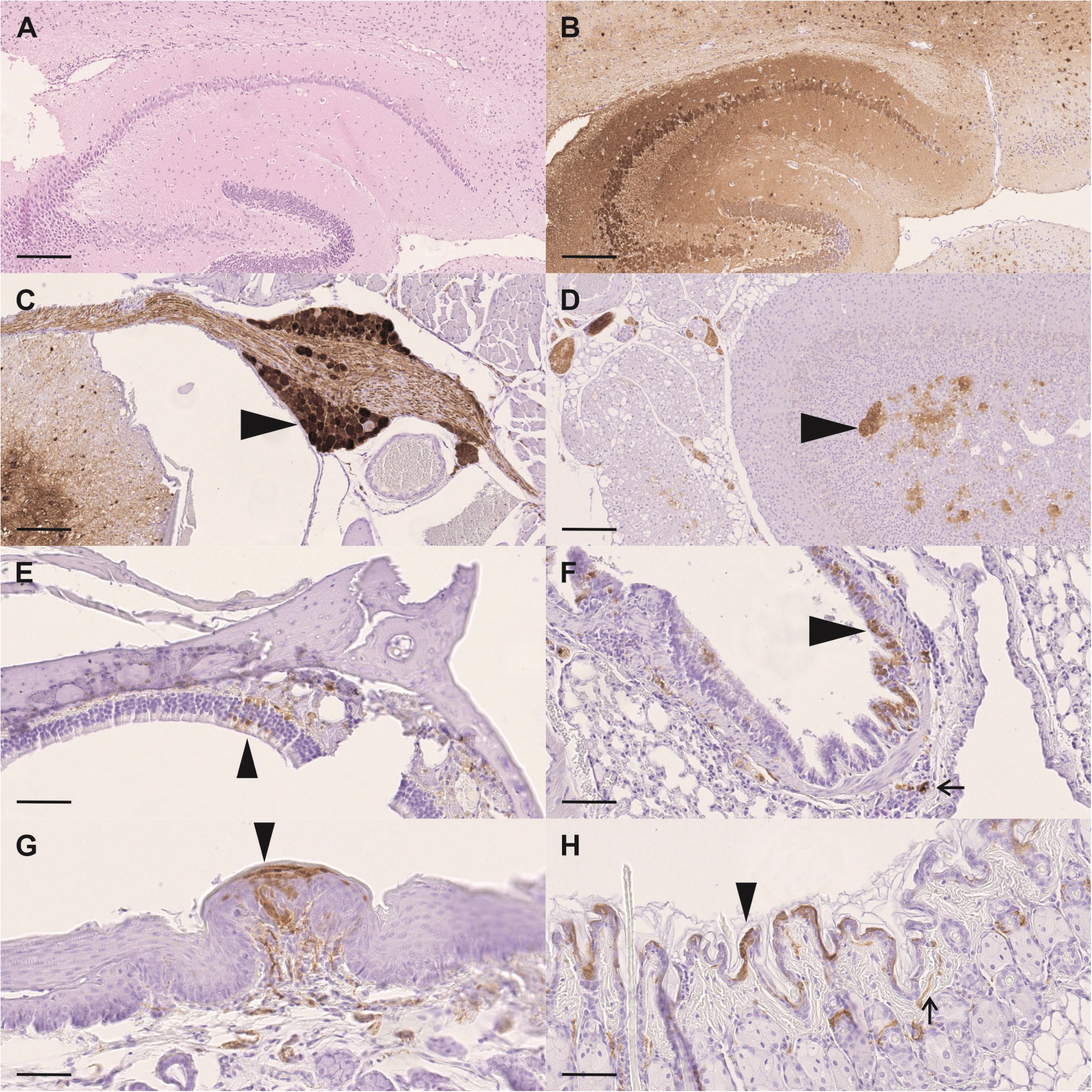
Absence of pathological lesions demonstrated with haematoxylin and eosin stain (A), and distribution of Borna disease virus 1 (BoDV1)-N-antigen demonstrated with immunohistochemistry depicted as brown signal (B-H) in shrews experimentally infected with BoDV1. A. Intracerebrally infected shrew (H43) with no inflammatory or degenerative lesions, scale bar = 250µm. B. Same animal as in A with abundant BoDV1-N antigen in neurons, astrocytes, and in the neuropil, scale bar = 250µm. C. Intracerebrally infected shrew (H49) displays BoDV1-N-antigen in spinal ganglion nuclei (arrowhead) and axons, scale bar = 100µm. D. Adrenal gland of a subcutaneously infected shrew (H41) with BoDV1-N-antigen in medullary adrenal cells (arrowhead), scale bar = 200µm. E. Evaluation of nasal cavity of intracerebrally infected shrew (H16) yields positive BoDV1-N-antigen in olfactory epithelial cells (arrowhead), scale bar = 100µm. F. The lung of an intranasally infected shrew (H53) displays BoDV1-N-antigen in both cytoplasm and nuclei of bronchial epithelium (arrowhead) and submucosal nerve fibres (arrow), scale bar = 100µm. G. Intraperitoneally infected shrew (H48) with BoDV1-N-antigen in epithelial cells of taste papillae (arrowhead), scale bar = 50µm. H. Skin of the same animal as in F with BoDV1-N-virus antigen in the cytoplasm of epidermal keratinocytes (arrowhead) and dermal nerve fibres (arrow), scale bar = 100µm. BoDV1, Borna disease virus 1.

**Table 3.**
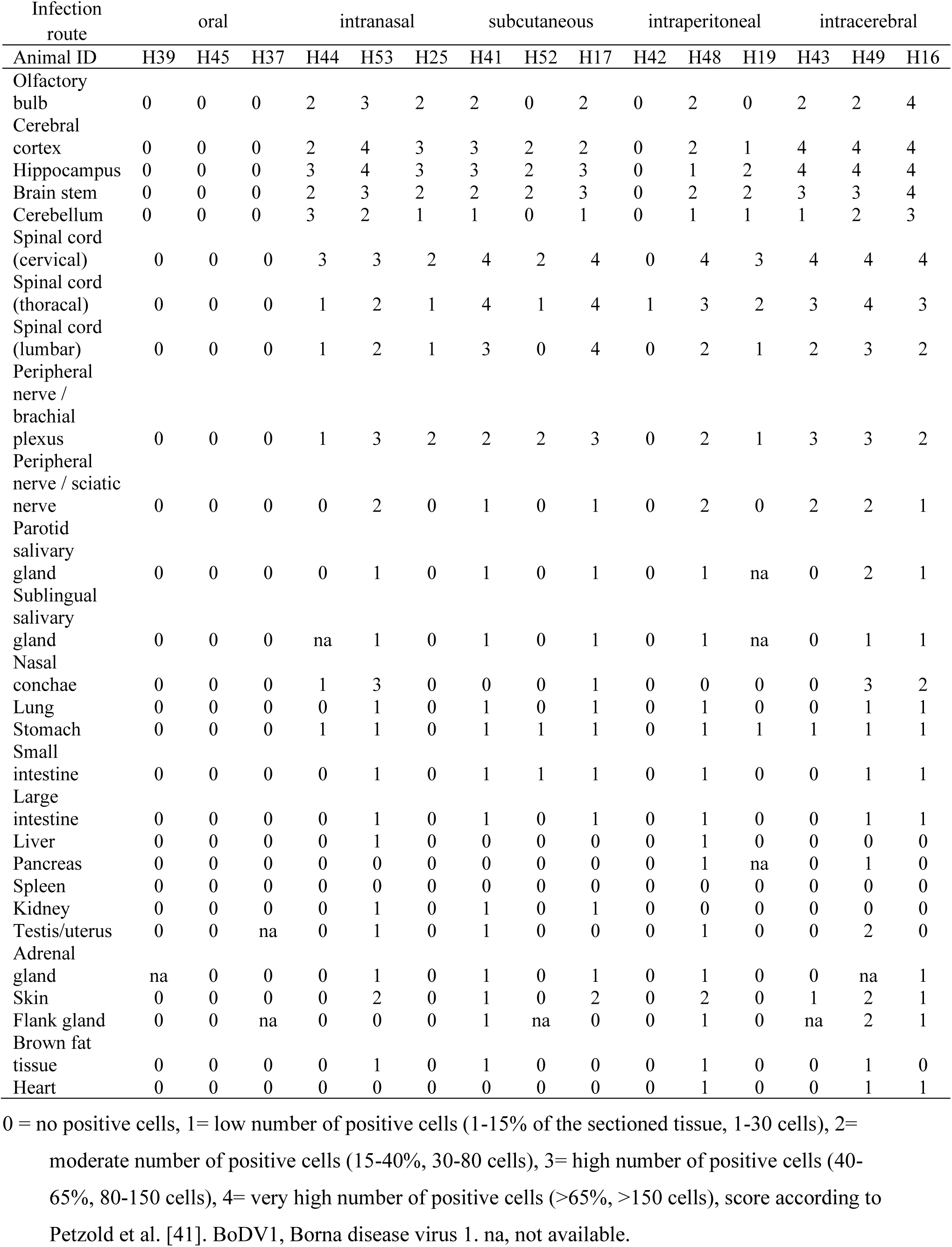
Demonstration of BoDV1 antigen distribution by immunohistochemistry.

| Infection route | oral |  |  | intranasal |  |  | subcutaneous |  |  | intraperitoneal |  |  | intracerebral |  |  |
| --- | --- | --- | --- | --- | --- | --- | --- | --- | --- | --- | --- | --- | --- | --- | --- |
| Animal ID | H39 | H45 | H37 | H44 | H53 | H25 | H41 | H52 | H17 | H42 | H48 | H19 | H43 | H49 | H16 |
| Olfactory bulb | 0 | 0 | 0 | 2 | 3 | 2 | 2 | 0 | 2 | 0 | 2 | 0 | 2 | 2 | 4 |
| Cerebral cortex | 0 | 0 | 0 | 2 | 4 | 3 | 3 | 2 | 2 | 0 | 2 | 1 | 4 | 4 | 4 |
| Hippocampus | 0 | 0 | 0 | 3 | 4 | 3 | 3 | 2 | 3 | 0 | 1 | 2 | 4 | 4 | 4 |
| Brain stem | 0 | 0 | 0 | 2 | 3 | 2 | 2 | 2 | 3 | 0 | 2 | 2 | 3 | 3 | 4 |
| Cerebellum | 0 | 0 | 0 | 3 | 2 | 1 | 1 | 0 | 1 | 0 | 1 | 1 | 1 | 2 | 3 |
| Spinal cord (cervical) | 0 | 0 | 0 | 3 | 3 | 2 | 4 | 2 | 4 | 0 | 4 | 3 | 4 | 4 | 4 |
| Spinal cord (thoracal) | 0 | 0 | 0 | 1 | 2 | 1 | 4 | 1 | 4 | 1 | 3 | 2 | 3 | 4 | 3 |
| Spinal cord (lumbar) | 0 | 0 | 0 | 1 | 2 | 1 | 3 | 0 | 4 | 0 | 2 | 1 | 2 | 3 | 2 |
| Peripheral nerve / brachial plexus | 0 | 0 | 0 | 1 | 3 | 2 | 2 | 2 | 3 | 0 | 2 | 1 | 3 | 3 | 2 |
| Peripheral nerve / sciatic nerve | 0 | 0 | 0 | 0 | 2 | 0 | 1 | 0 | 1 | 0 | 2 | 0 | 2 | 2 | 1 |
| Parotid salivary gland | 0 | 0 | 0 | 0 | 1 | 0 | 1 | 0 | 1 | 0 | 1 | na | 0 | 2 | 1 |
| Sublingual salivary gland | 0 | 0 | 0 | na | 1 | 0 | 1 | 0 | 1 | 0 | 1 | na | 0 | 1 | 1 |
| Nasal conchae | 0 | 0 | 0 | 1 | 3 | 0 | 0 | 0 | 1 | 0 | 0 | 0 | 0 | 3 | 2 |
| Lung | 0 | 0 | 0 | 0 | 1 | 0 | 1 | 0 | 1 | 0 | 1 | 0 | 0 | 1 | 1 |
| Stomach | 0 | 0 | 0 | 1 | 1 | 0 | 1 | 1 | 1 | 0 | 1 | 1 | 1 | 1 | 1 |
| Small intestine | 0 | 0 | 0 | 0 | 1 | 0 | 1 | 1 | 1 | 0 | 1 | 0 | 0 | 1 | 1 |
| Large intestine | 0 | 0 | 0 | 0 | 1 | 0 | 1 | 0 | 1 | 0 | 1 | 0 | 0 | 1 | 1 |
| Liver | 0 | 0 | 0 | 0 | 1 | 0 | 0 | 0 | 0 | 0 | 1 | 0 | 0 | 0 | 0 |
| Pancreas | 0 | 0 | 0 | 0 | 0 | 0 | 0 | 0 | 0 | 0 | 1 | na | 0 | 1 | 0 |
| Spleen | 0 | 0 | 0 | 0 | 0 | 0 | 0 | 0 | 0 | 0 | 0 | 0 | 0 | 0 | 0 |
| Kidney | 0 | 0 | 0 | 0 | 1 | 0 | 1 | 0 | 1 | 0 | 0 | 0 | 0 | 0 | 0 |
| Testis/uterus | 0 | 0 | na | 0 | 1 | 0 | 1 | 0 | 0 | 0 | 1 | 0 | 0 | 2 | 0 |
| Adrenal gland | na | 0 | 0 | 0 | 1 | 0 | 1 | 0 | 1 | 0 | 1 | 0 | 0 | na | 1 |
| Skin | 0 | 0 | 0 | 0 | 2 | 0 | 1 | 0 | 2 | 0 | 2 | 0 | 1 | 2 | 1 |
| Flank gland | 0 | 0 | na | 0 | 0 | 0 | 1 | na | 0 | 0 | 1 | 0 | na | 2 | 1 |
| Brown fat tissue | 0 | 0 | 0 | 0 | 1 | 0 | 1 | 0 | 0 | 0 | 1 | 0 | 0 | 1 | 0 |
| Heart | 0 | 0 | 0 | 0 | 0 | 0 | 0 | 0 | 0 | 0 | 1 | 0 | 0 | 1 | 1 |

0 = no positive cells, 1= low number of positive cells (1-15% of the sectioned tissue, 1-30 cells), 2= moderate number of positive cells (15-40%, 30-80 cells), 3= high number of positive cells (40-65%, 80-150 cells), 4= very high number of positive cells (>65%, >150 cells), score according to Petzold et al. [41]. BoDV1, Borna disease virus 1. na, not available.

In the spinal cord, all intranasally infected animals, as well as each two of the intraperitoneally, intracerebrally and subcutaneously infected animals displayed decreasing BoDV1-N distribution and number of positive cells from the cervical to the lumbar spinal cord. One subcutaneously and one intracranially infected animal displayed comparable distribution and number of positive cells throughout the whole spinal cord. Both grey and white matter were affected in all those animals, including motor neurons, glia cells and surrounding neuropil with cellular processes. Additionally, dorsal root ganglia, and if evaluable, nerve fibres within the surrounding skeletal muscle, were consistently positive. Notably, a mediodorsal triangular area within the dorsal funiculus of the spinal cord was relatively spared in evaluation of BoDV1-N-antigen distribution, displaying consistently less or no positive fibres (Figure 5).

**Figure 5.**
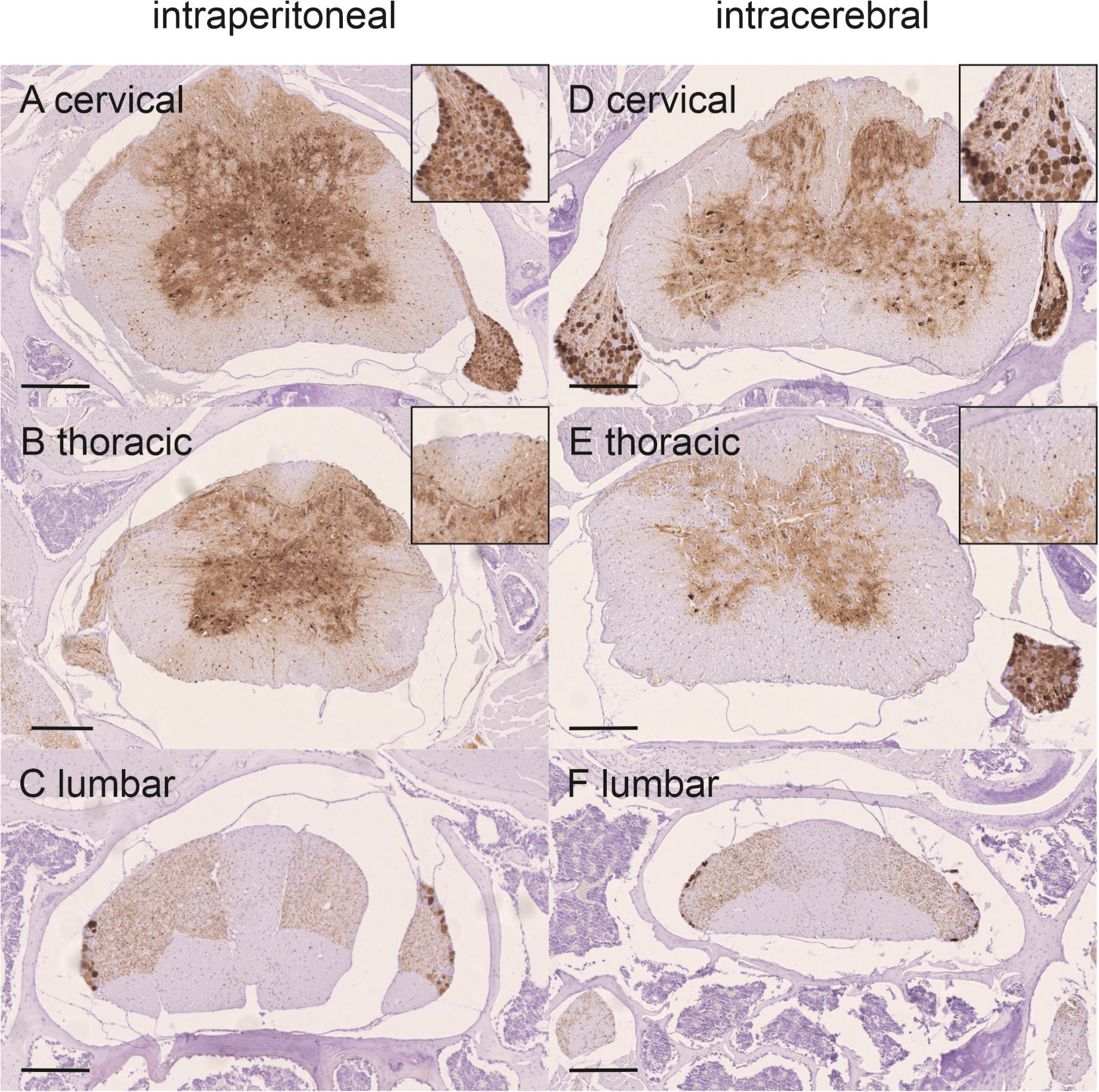
Overview of Borna disease virus 1 (BoDV1)-N antigen distribution in the spinal cord of experimentally BoDV1 infected shrews by intraperitoneal infection (A-C) and intracerebral infection (D-E) demonstrated with immunohistochemistry depicted as brown signal. A. Cervical spinal cord of an intraperitoneally infected shrew with BoDV1-N-antigen in both grey and white matter and including motor neurons, glia cells and surrounding neuropil as well as dorsal root ganglia (inlay), scale bar = 500 µm. B. Thoracic spinal cord of an intraperitoneally infected shrew with BoDV1-N-antigen in both grey and white matter and including motor neurons, glia cells and surrounding neuropil as well as dorsal root ganglia. A mediodorsal triangular area within the dorsal funiculus of the spinal cord corresponding histologically to the gracile fasciculus displays hardly positive fibres (inlay), scale bar = 250 µm. C. Lumbar spinal cord of an intraperitoneally infected shrew with BoDV1-N-antigen in both grey and white matter and including motor neurons, glia cells and surrounding neuropil, scale bar = 250 µm. D. Cervical spinal cord of an intracerebrally infected shrew with BoDV1-N-antigen in both grey and white matter and including motor neurons, glia cells and surrounding neuropil as well as dorsal root ganglia (inlay), scale bar = 500 µm. E. Thoracic spinal cord of an intracerebrally infected shrew with BoDV1-N-antigen in both grey and white matter and including motor neurons, glia cells and surrounding neuropil as well as dorsal root ganglia. A mediodorsal triangular area within the dorsal funiculus of the spinal cord corresponding histologically to the gracile fasciculus displays hardly positive fibres (inlay), scale bar = 250 µm. F. Lumbar spinal cord of an intracerebrally infected shrew with BoDV1-N-antigen in both grey and white matter and including motor neurons, glia cells and surrounding neuropil, scale bar = 250 µm.

In one intraperitoneally infected animal BoDV1-N-antigen was not detected in any section and part of the spinal cord, except for a centrally located neuron and its immediate surrounding neuropil within the thoracic grey matter, and unilaterally, very few nerve fibres ventrally of and surrounding the thoracic vertebral column. Peripheral nerve fibres in parotid and sublingual salivary gland, lung, kidney, skin, uterus, testis, and heart displayed BoDV1-N-antigen, as well as myenteric ganglia in the stomach and intestine.

Besides nervous tissue, epithelial and mesenchymal cells occasionally displayed BoDV1-antigen both intracytoplasmically and intranuclearly in all infected cohorts. Occasionally, epithelial cells of the parotid and sublingual salivary gland displayed viral antigen, as well as epithelial cells of the lingual taste papillae. Besides nerve fibres of the skin and lung, antigen was also found in epidermal keratinocytes and bronchial epithelial cells, respectively.

Additionally, some pancreatic islets cells and mesenchymal cells of the brown fat tissue harboured BoDV1-N antigen. There was no perceptible difference between the infection groups in these above-mentioned tissues. In the nasal conchae, viral antigen was observed in nerve fibres and olfactory epithelial cells of the intracerebrally and intranasally infected animals, whereas subcutaneously infected animals displayed antigen solely in nerve fibres.

## Discussion

Emerging and re-emerging diseases caused by pathogens originating from wildlife reservoirs is of major concern for public health worldwide [1,2]. Next to human health, One-Health approaches also put a spotlight on the environment, as well as on domestic animals and wildlife in their natural habitats [24,25]. Due to the availability of various high-throughput methods, such as metagenomics, our knowledge of potential pathogens, as well as spillover and reservoir host species is constantly expanding [1,42] and may potentially extent to include additional currently unknown animal, human and environmental factors.

In Germany, increasing numbers of human encephalitis cases caused by infection with BoDV1 have raised the awareness und urgent need to understand underlying pathogen-reservoir host interactions and potential spillover events [4,6,23]. Besides sampling of naturally infected shrew species, detailed studies were hampered due to the lack of suitable animal- and *in vitro* models to investigate the reservoir situation. On the contrary, models for the analysis of the dead-end host status, with viral restriction to the CNS and concomitant non-purulent meningoencephalitis based on T cell-mediated immunopathology, have been widely used and well characterized to date. Furthermore, neonatal rats have already been successfully used to investigate a potential immune tolerance to BoDV1 infections and showed remarkable concordance of infection results compared to the reservoir *C. leucodon* [10,17,18,20,39]. To further expand knowledge on the BoDV1 infection of the reservoir, *C. russula*, the greater white-toothed shrew, was examined for its suitability as a shrew model to study the biology of bornavirus infections and their underlying pathogenesis in the reservoir. As the husbandry and breeding of *C. russula* has already been successfully established for more than two years, a laboratory shrew colony from comparable husbandry conditions was available in our working group.

In general, experimental BoDV1 infection of greater white-toothed shrews was successful by various routes such as intranasal, subcutaneous, intraperitoneal, and intracerebral inoculation, indicating its usefulness as animal model species. More importantly, our results support the possibility for *C. russula* to function as a natural wildlife reservoir for BoDV1, with potential of spillover to other species due to viral shedding. This observation has recently been supported by the detection of viral RNA in tissue pools from one *C. russula* individual from Germany [30]. However, in comparison to the vast number of BoDV1-positive *C. leucodon* detected over time [17–19,29,30], this represents a rather rare but nonetheless possible event. The successful and simpler husbandry and breeding of *C. russula*, and its hereby proven susceptibility to BoDV1 infection make this shrew species a suitable laboratory shrew model.

The mode of virus transmission within the shrew population as well as to dead end hosts, such as horses and humans, is still under investigation. Former studies [18,19] suggested the olfactory epithelium as the most likely entry site for BoDV1 infection in shrews, because it is also a plausible entry site for spillover hosts such as horses [43]. Successful intranasal infection of rats in former studies [43,44], and of greater white-toothed shrews in this study, further support this hypothesis. However, since subcutaneous and intraperitoneal infections were also successful in *C. russula*, other or additional potential entry sites, e.g., via skin lesions and wounds, might be possible. This has also already been shown for avian bornaviruses [45]. Since oral inoculation did not cause BoDV1 infection, this route of entry does not seem as likely as the other investigated inoculation routes in shrews, comparable to the transmission of avian bornaviruses [45,46].

The presence of infectious BoDV1 in saliva, and BoDV1 RNA in skin swabs and faeces at 35 days p.i. suggests start of shedding by saliva, skin, and faeces, similarly to *C. leucodon* [20] and corresponding to demonstration of BoDV1 antigen in epithelial cells of salivary glands and epidermal keratinocytes. A previous study, performed on persistently BoDV1 infected bicolored white-toothed shrews by our research group, already demonstrated that *C. leucodon* shed infectious virus and RNA for a long period through saliva, skin, lacrimal fluid, urine, and faeces [20]. However, the shedding of BoDV1 in this study was inconsistent over the observed time course of 41 days, and therefore other, yet unknown, factors may also influence the extent of viral shedding. In BoDV1-infected *C. russula*, viral shedding could not be detected in all animals at the end point of the experiment, which could indicate that virus shedding starts around 35 to 41 days p.i. Differences might also be due to route of infection determining the time frame for the virus to reach the peripheral organs. In summary, these data suggests that viral spread to organs capable of secretion and excretion may require about or more than six weeks after BoDV1 infection and may also depend on the initial entry site of the virus. Intracerebrally BoDV1-infected animals were those that already shed infectious virus via saliva on day 41 p.i., while in all other groups solely viral RNA was detected in the swabs. In combination with the highest viral load in the CNS (brain and spinal cord) this argues for viral spread from the CNS to the periphery in case of an intracerebral infection. In case of peripheral inoculation routes, the spinal cord also harbours high viral loads, concluding that spread from periphery to the CNS to enhance viral replication, and subsequent centrifugal spread to organs capable of secretion and excretions seems feasible.

Notably, a mediodorsal triangular area within the dorsal funiculus of the spinal cord was relatively spared in evaluation of BoDV1-N-antigen distribution, displaying consistently less or no positive fibres (Figure 5). No tracing studies have been done to characterize spinal tracts and their exact pathways and projections in *Crocidura* sp., but the area corresponds histologically to the gracile fasciculus and/or including the postsynaptic pathways of the dorsal column in mice and rats. In correlation with infection route, this might indicate that ascending sensory nerve fibres from the distal limbs are less frequently or less extensively used for BoDV1 trafficking. However, frequent positive signals in dorsal and ventral roots, extraspinal nerve fibres, ganglia ventrally to the spinal cord, as well as consistent signals in dorsal root ganglia, suggest that BoDV1 may travel along at both sensory and motor neuron axons. Nevertheless, in-depth studies of the crocidurine nervous system are needed to elucidate neural pathways, their functional aspects, and to exclude differences in fibre density in the dorsal funiculus as cause for this particular observation.

In BoDV1-infected *C. russula*, viral antigen was already present in a few clusters of epithelial cells, which might represent a prerequisite for later viral shedding. Retrograde spread from peripheral inoculation sites into the CNS, with centrifugal spread to the periphery has already been demonstrated after peripheral inoculation of avian bornaviruses in cockatiels [47]. Only few epithelial cells were BoDV1-N-antigen-positive in this study with *C. russula*, compared to former studies with *C. leucodon* [18,19,29], most likely indicating the beginning of viral infection of the periphery. Viral shedding increased at the end of the here presented study, concluding that a longer period of infection might be needed to establish viral spread to all organs, and subsequent viral shedding via various routes in *C. russula*.

In contrast to persistently BoDV1 infected *C. leucodon* with an asymptomatic course of chronic infection [20], the experimental infection of *C. russula* led to reduced feed intake and weight loss in the acute phase. In the first week after infection, this could also be associated with anaesthesia performed to conduct experimental BoDV1 inoculation, as orally, but unsuccessful inoculation similarly led to reduced feed intake. Weight loss and reduced feed intake was not associated with any morphological lesion in the CNS, or any other organ system of the animals euthanized before 41 dpi. Further studies should monitor more physiological parameters, such as body temperature or heart rate, to detect potential deviations. Furthermore, it is difficult to assess whether the observed reduced feed intake and weight loss might represent typical features of the acute phase of natural BoDV1 infections in shrews. Weight loss and starvation can enhance death, and/or affect the detection of infected animals by predators. However, it is questionable whether this might explain the small numbers of naturally BoDV1-infected *C. russula* detected so far, as more factors are most likely involved. Whether any other conditions of the experimental infection, such as virus dose, affect the course of infection and facilitate the observed clinical signs, also needs to be addressed in future studies.

Interestingly, the absence of organ lesions, despite viral replication and spread in infected *C. russula*, points to a comparable virus tolerance and establishment of viral persistence as seen in *C. leucodon*. This further substantiates *C. russula* as a suitable shrew BoDV1-infection model. The reason for the absence or evasion of an antiviral immune response and virus elimination is not yet known for the natural shrew reservoir. As Lewis rats develop a comparable pattern of virus distribution and shedding without organ lesions if they are infected as immune-incompetent newborns [39], some kind of immune tolerance seems likely in the shrew. However, since *C. russula* in this study were successfully infected as adults, different and so far, unknown mechanisms of immune tolerance might be operative in the shrew. Infected *C. russula* developed high serum antibody titres, comparable to the neonatally infected Lewis rats [39]. This further substantiates that the humoral antiviral response does not protect against extended viral spread and shedding, mediated by virus-specific antibodies.

In summary, the successful experimental infection of *C. russula* supports their role as suitable shrew model and strengthens the status of crocidurine shrews as reservoir hosts of BoDV1 in endemic areas. The possibility of different successful infection routes suggests that several transmission pathways are possible to maintain the virus within shrew populations. The onset of virus shedding after 5-6 weeks p.i. is indicative of the period of infection required to become an exposure risk to susceptible dead-end hosts such as humans or horses. To date, further questions remain unanswered, particularly regarding the potential immune tolerance of the shrews, the route of transmission to dead end hosts, the long-term maintenance of BoDV1 infection in reservoir populations, and the various underlying pathogenetic mechanisms. As a prospect, this novel and successfully established shrew model can open the avenue to address these questions in future studies.

## Acknowledgements

This work was supported by grants from the Federal Ministry of Education and Research project ZooBoCo (Bundesministerium für Bildung und Forschung) (Zoonotic Bornavirus Consortium), within the research network for zoonotic infectious diseases, grant number 01KI1722E/ 01KI2005E.

## Disclosure Statement

The authors report there are no competing interests to declare.

## Data Availability Statement

The data that support the findings of this study are available from the corresponding authors, (DN, CH), upon reasonable request.

## Supplementary data

**Supplemental table 1.** Score Sheet: Evaluation of the well-being of *Crocidura russula*.

|  |  |
| --- | --- |
| 1. Daily food intake |  |
| >0.8x of body mass | 0 |
| >0.7x and >0.79x of body mass | 1 |
| >0.5x and >0.69x of body mass | 2 |
| <0.5x of body mass | 3 |
| 2. Physical appearance |  |
| Coat smooth and shining, orifices of the body clean | 0 |
| Dull coat | 1 |
| Coat staring, ocular and nasal discharges, anus daubed | 2 |
| Piloerection, open wounds, closed eyes | 3 |
| 3. Natural behaviour |  |
| Attentive, curious, swift movements, foraging or hiding | 0 |
| Inattentive, reduced and slow or excessive movement, unexplained behaviour | 1 |
| reduced movement, obvious movement pain | 2 |
| Unconscious, no movement, weak, trembling, heavy breathing | 3 |
| 4. Breathing |  |
| normal regular and relaxed breathing | 0 |
| mildly increased breathing | 1 |
| moderate increased breathing, abdominal breathing | 2 |
| heavy breathing and gasping | 3 |
| 5. Body Mass |  |
| Compared to body mass measured less than 14 days before |  |
| Identical or increased | 0 |
| 5%-10% reduction | 1 |
| 10 % - 20% reduction | 2 |
| Reduction of >20 % | 3 |
| Interpretation<br>Score of 0-1, Characteristics of healthy shrews<br>Score of 2-3, Signs of mild suffering<br>Score of 4-7, Signs of moderate suffering, consultation of veterinarian, increase monitoring, euthanasia by score of 4-5 during 3 days or score of 6-7 during 3 days<br>Score of 8, Signs of severe suffering, euthanasia |  |

**Supplementary table 2.** Overview of Cq values (respective left column) and viral copy numbers (respective right column) of evaluated tissues from each animal

| Infection route |  |  | oral |  |  |  |
| --- | --- | --- | --- | --- | --- | --- |
| Animal ID |  | H39 |  | H45 |  | H37 |
| Olfactory bulb | neg | <1000 | neg | <1000 | neg | <1000 |
| Cerebral cortex | neg | <1000 | neg | <1000 | neg | <1000 |
| Hippocampus | neg | <1000 | neg | <1000 | neg | <1000 |
| Brain stem | neg | <1000 | neg | <1000 | neg | <1000 |
| Cerebellum | neg | <1000 | neg | <1000 | neg | <1000 |
| Peripheral nerve /<br>brachial plexus | neg | <1000 | neg | <1000 | neg | <1000 |
| Peripheral nerve / sciatic<br>nerve | neg | <1000 | neg | <1000 | neg | <1000 |
| Parotid salivary gland | neg | <1000 | neg | <1000 | neg | <1000 |
| Sublingual salivary gland | neg | <1000 | neg | <1000 | neg | <1000 |
| Nasal conchae | neg | <1000 | neg | <1000 | neg | <1000 |
| Lung | neg | <1000 | neg | <1000 | neg | <1000 |
| Oesophagus | neg | <1000 | neg | <1000 | neg | <1000 |
| Stomach | neg | <1000 | neg | <1000 | neg | <1000 |
| Small intestine | neg | <1000 | neg | <1000 | neg | <1000 |
| Large intestine | neg | <1000 | neg | <1000 | neg | <1000 |
| Liver | neg | <1000 | neg | <1000 | neg | <1000 |
| Spleen | neg | <1000 | neg | <1000 | neg | <1000 |
| Bone marrow | neg | <1000 | neg | <1000 | neg | <1000 |
| Kidney | neg | <1000 | neg | <1000 | neg | <1000 |
| Urinary bladder | na | na | neg | <1000 | neg | <1000 |
| Testis/uterus | neg | <1000 | neg | <1000 | neg | <1000 |
| Adrenal gland | neg | <1000 | neg | <1000 | neg | <1000 |
| Skin | neg | <1000 | neg | <1000 | neg | <1000 |
| Flank gland | neg | <1000 | neg | <1000 | neg | <1000 |
| Skeletal muscle | neg | <1000 | neg | <1000 | neg | <1000 |
| Heart | neg | <1000 | neg | <1000 | neg | <1000 |

| Infection route |  | intranasal |  |  |  |  |
| --- | --- | --- | --- | --- | --- | --- |
| Animal ID |  | H44 |  | H53 |  | H25 |
| Olfactory bulb | 19,66 | 1,76E+09 | 16,74 | 3,79E+09 | 27,41 | 1,12E+07 |
| Cerebral cortex | 21,98 | 3,82E+08 | 19,1 | 8,54E+08 | 21,15 | 6,62E+08 |
| Hippocampus | 14,55 | 1,85E+09 | 15,19 | 1,01E+10 | 20,34 | 1,00E+09 |
| Brain stem | 20,34 | 1,12E+09 | 16,05 | 5,83E+09 | 20,55 | 9,68E+08 |
| Cerebellum | 22,25 | 3,20E+08 | 16,54 | 4,28E+09 | 27,45 | 1,09E+07 |
| Peripheral nerve /<br>brachial plexus | 30,27 | 1,82E+06 | 24,88 | 2,22E+07 | 30,61 | 1,38E+06 |
| Peripheral nerve / sciatic<br>nerve | 34,93 | 1,06E+05 | 27,65 | 3,88E+06 | 32 | 6,58E+05 |
| Parotid salivary gland | 33,16 | 2,80E+05 | 25,89 | 1,17E+07 | 29,41 | 3,10E+06 |
| Sublingual salivary gland | 28,53 | 5,40E+06 | 21 | 2,56E+08 | neg | <1000 |
| Nasal conchae | 22,97 | 2,02E+08 | 18,43 | 1,30E+09 | 32,99 | 4,10E+05 |
| Lung | neg | <1000 | 26,46 | 8,21E+06 | neg | <1000 |
| Oesophagus | 33,37 | 2,52E+05 | 25,81 | 1,24E+07 | 29,02 | 4,06E+06 |
| Stomach | 33,59 | 2,34E+05 | 23,71 | 4,64E+07 | neg | <1000 |
| Small intestine | neg | <1000 | 25,91 | 1,16E+07 | neg | <1000 |
| Large intestine | neg | <1000 | 26,43 | 8,37E+06 | neg | <1000 |
| Liver | neg | <1000 | neg | <1000 | neg | <1000 |
| Spleen | neg | <1000 | 31,08 | 4,45E+05 | neg | <1000 |
| Bone marrow | neg | <1000 | 28,18 | 2,78E+06 | 30,03 | 2,04E+06 |
| Kidney | neg | <1000 | 29,21 | 1,45E+06 | neg | <1000 |
| Urinary bladder | neg | <1000 | 30,89 | 5,02E+05 | neg | <1000 |
| Testis/uterus | 27,46 | 1,09E+07 | neg | <1000 | neg | <1000 |
| Adrenal gland | neg | <1000 | 25,31 | 1,69E+07 | neg | <1000 |
| Skin | 30,81 | 1,34E+06 | 27,95 | 3,20E+06 | 31,66 | 7,36E+05 |
| Flank gland | neg | <1000 | 27,72 | 3,71E+06 | neg | <1000 |
| Skeletal muscle | neg | <1000 | 30,36 | 7,02E+05 | 33,43 | 2,74E+05 |
| Heart | neg | <1000 | 29,63 | 1,11E+06 | neg | <1000 |

| Infection route | subcutaneous |  |  |  |  |  |
| --- | --- | --- | --- | --- | --- | --- |
| Animal ID | H41 |  | H52 |  | H17 |  |
| Olfactory bulb | 23,46 | 5,69E+07 | 21,13 | 1,44E+08 | 22,77 | 7,64E+07 |
| Cerebral cortex | 21,05 | 1,41E+08 | 22,14 | 7,26E+07 | 19,12 | 3,65E+08 |
| Hippocampus | 19,9 | 2,17E+08 | 21,12 | 2,14E+07 | 17,99 | 5,93E+08 |
| Brain stem | 17,75 | 4,89E+08 | 22,17 | 7,12E+07 | 17,19 | 8,36E+08 |
| Cerebellum | 19,77 | 2,28E+08 | 27,97 | 1,41E+06 | 19,13 | 3,64E+08 |
| Peripheral nerve /<br>brachial plexus | 24,54 | 3,78E+07 | 31,2 | 1,59E+05 | 22,51 | 8,54E+07 |
| Peripheral nerve / sciatic<br>nerve | 27,58 | 1,20E+07 | 29,65 | 4,52E+05 | 28,89 | 5,58E+06 |
| Parotid salivary gland | 30,45 | 4,07E+06 | neg | <1000 | 23,96 | 4,59E+07 |
| Sublingual salivary gland | 31,75 | 2,50E+06 | 24,78 | 1,22E+07 | neg | <1000 |
| Nasal conchae | 27,29 | 1,34E+07 | 30,77 | 2,12E+05 | 25,83 | 2,06E+07 |
| Lung | neg | <1000 | 33,55 | 3,22E+04 | neg | <1000 |
| Oesophagus | 23,71 | 5,16E+07 | 33,04 | 4,56E+04 | 23,17 | 6,46E+07 |
| Stomach | 27,72 | 1,14E+07 | 27,78 | 1,60E+06 | 26,02 | 1,90E+07 |
| Small intestine | 26,47 | 1,83E+07 | 31,15 | 1,64E+05 | 29,23 | 4,82E+06 |
| Large intestine | 28,3 | 9,18E+06 | 29,4 | 5,36E+05 | 26,55 | 1,52E+07 |
| Liver | neg | <1000 | neg | <1000 | neg | <1000 |
| Spleen | 31,95 | 2,32E+06 | 32,36 | 7,22E+04 | neg | <1000 |
| Bone marrow | 32,61 | 1,81E+06 | 29,59 | 4,70E+05 | 27,61 | 9,61E+06 |
| Kidney | 29,07 | 6,87E+06 | 27,88 | 1,49E+06 | 31,97 | 1,49E+06 |
| Urinary bladder | 32,43 | 1,94E+06 | 29,24 | 5,96E+05 | 26,8 | 1,36E+07 |
| Testis/uterus | 25,28 | 2,86E+07 | 31,84 | 1,03E+05 | neg | <1000 |
| Adrenal gland | 21 | 1,43E+08 | 26,15 | 4,82E+06 | 23,7 | 5,15E+07 |
| Skin | 23,49 | 5,62E+07 | 22,06 | 7,68E+07 | 23,59 | 5,38E+07 |
| Flank gland | 23,34 | 5,95E+07 | 29,15 | 6,34E+05 | 26,43 | 1,60E+07 |
| Skeletal muscle | 28,41 | 8,78E+06 | 30,98 | 1,84E+05 | 25,31 | 2,58E+07 |
| Heart | 31,04 | 3,27E+06 | neg | <1000 | neg | <1000 |

| Infection route |  | intraperitoneal |  |  |  |  |
| --- | --- | --- | --- | --- | --- | --- |
| Animal ID |  | H42 | H48 |  | H19 |  |
| Olfactory bulb | neg | <1000 | 24,11 | 4,30E+07 | neg | <1000 |
| Cerebral cortex | neg | <1000 | 21,37 | 1,39E+08 | 27,56 | 1,86E+06 |
| Hippocampus | neg | <1000 | 18,8 | 4,19E+08 | 23,04 | 7,28E+07 |
| Brain stem | neg | <1000 | 18,01 | 5,88E+08 | 24,86 | 1,15E+07 |
| Cerebellum | neg | <1000 | 18,83 | 4,13E+08 | neg | <1000 |
| Peripheral nerve /<br>brachial plexus | neg | <1000 | 19,76 | 2,77E+08 | 23,75 | 2,44E+07 |
| Peripheral nerve / sciatic<br>nerve | neg | <1000 | 24,7 | 3,35E+07 | 30,83 | 2,02E+05 |
| Parotid salivary gland | neg | <1000 | 22,75 | 7,71E+07 | 31,86 | 1,02E+05 |
| Sublingual salivary gland | neg | <1000 | 33,38 | 8,15E+05 | 27,9 | 1,48E+06 |
| Nasal conchae | neg | <1000 | 30,25 | 3,11E+06 | 32,9 | 5,02E+04 |
| Lung | neg | <1000 | 34,6 | 4,84E+05 | 33,49 | 3,36E+04 |
| Oesophagus | neg | <1000 | 25,51 | 2,37E+07 | 28,25 | 1,16E+06 |
| Stomach | neg | <1000 | 28,18 | 7,54E+06 | 34,73 | 1,46E+04 |
| Small intestine | neg | <1000 | 22,32 | 9,27E+07 | 32,72 | 5,68E+04 |
| Large intestine | neg | <1000 | 22,13 | 1,00E+08 | 30,54 | 2,46E+05 |
| Liver | neg | <1000 | neg | <1000 | neg | <1000 |
| Spleen | neg | <1000 | 28,34 | 7,04E+06 | neg | <1000 |
| Bone marrow | neg | <1000 | 29,87 | 3,65E+06 | 27,58 | 1,83E+06 |
| Kidney | neg | <1000 | 31,98 | 1,48E+06 | neg | <1000 |
| Urinary bladder | neg | <1000 | 29,38 | 4,52E+06 | neg | <1000 |
| Testis/uterus | neg | <1000 | neg | <1000 | 27,34 | 2,16E+06 |
| Adrenal gland | neg | <1000 | 21,49 | 1,32E+08 | neg | <1000 |
| Skin | neg | <1000 | 22,39 | 9,01E+07 | 28,82 | 7,94E+05 |
| Flank gland | neg | <1000 | 30,22 | 3,15E+06 | 28,78 | 8,16E+05 |
| Skeletal muscle | neg | <1000 | 28,24 | 7,36E+06 | 31 | 1,82E+05 |
| Heart | neg | <1000 | 30,66 | 2,61E+06 | neg | <1000 |

| Infection route | intracerebral |  |  |  |  |  |
| --- | --- | --- | --- | --- | --- | --- |
| Animal ID | H43 |  | H49 |  | H16 |  |
| Olfactory bulb | 17,34 | 2,22E+09 | 17,16 | 7,50E+09 | 15,28 | 4,03E+09 |
| Cerebral cortex | 15,34 | 8,45E+09 | 15,47 | 1,89E+10 | 14,99 | 4,72E+09 |
| Hippocampus | 14,09 | 1,96E+10 | 15,5 | 1,85E+10 | 13,51 | 1,06E+10 |
| Brain stem | 17,34 | 2,21E+09 | 16,09 | 1,35E+10 | 14,3 | 6,87E+09 |
| Cerebellum | 19,19 | 6,41E+08 | 18,86 | 2,97E+09 | 16,44 | 2,14E+09 |
| Peripheral nerve /<br>brachial plexus | 22,07 | 9,23E+07 | 23,06 | 3,00E+08 | 21,28 | 1,55E+08 |
| Peripheral nerve / sciatic<br>nerve | 23,86 | 2,79E+07 | 26,7 | 4,13E+07 | 25,3 | 1,74E+07 |
| Parotid salivary gland | 24,91 | 1,37E+07 | 24,92 | 1,09E+08 | 28,59 | 2,91E+06 |
| Sublingual salivary gland | 31,3 | 1,90E+05 | 27,77 | 2,30E+07 | 26,57 | 8,73E+06 |
| Nasal conchae | 23,21 | 4,31E+07 | 22,59 | 3,88E+08 | 18,22 | 8,16E+08 |
| Lung | 31,72 | 1,43E+05 | 28,62 | 1,45E+07 | 24,69 | 2,42E+07 |
| Oesophagus | 24,63 | 1,66E+07 | 25,24 | 9,12E+07 | 22,31 | 8,82E+07 |
| Stomach | 26,05 | 6,43E+06 | 24,76 | 1,19E+08 | 21,42 | 1,43E+08 |
| Small intestine | 30,61 | 3,01E+05 | 25,63 | 7,40E+07 | 21,48 | 1,39E+08 |
| Large intestine | 33,48 | 4,38E+04 | 29,1 | 1,11E+07 | 22,85 | 6,60E+07 |
| Liver | neg | <1000 | neg | <1000 | neg | <1000 |
| Spleen | neg | <1000 | neg | <1000 | 30,13 | 1,26E+06 |
| Bone marrow | 21,7 | 1,19E+08 | 26,7 | 4,11E+07 | 25,4 | 1,65E+07 |
| Kidney | neg | <1000 | neg | <1000 | 25,77 | 1,35E+07 |
| Urinary bladder | 33,95 | 3,20E+04 | 25,95 | 6,19E+07 | 24,86 | 2,20E+07 |
| Testis/uterus | 30,21 | 3,93E+05 | 26,62 | 4,29E+07 | neg | <1000 |
| Adrenal gland | 23,69 | 3,13E+07 | 25,74 | 6,96E+07 | 22,19 | 9,42E+07 |
| Skin | 26,14 | 6,03E+06 | 24,66 | 1,25E+08 | 24,17 | 3,22E+07 |
| Flank gland | 27,35 | 2,68E+06 | 26,25 | 5,27E+07 | 23,79 | 3,95E+07 |
| Skeletal muscle | 24,43 | 1,90E+07 | 26,71 | 4,09E+07 | 26,35 | 9,82E+06 |
| Heart | neg | <1000 | 34,15 | 7,06E+05 | 31,61 | 5,63E+05 |

Na, not available. Neg, negative (equals Cq above cycle 35 with viral copy numbers below the lower limit of 1000 viral copies)

